# Two-pore channel protein 2-mediated calcium release promotes angiopoietin 2 secretion by regulating Rab46-dependent Weibel-Palade body trafficking

**DOI:** 10.1101/2025.03.03.641167

**Authors:** Ryan D. Murray, Melissa Rose, Katarina T. Miteva, David J. Beech, Lynn McKeown

## Abstract

Angiopoietin2 (Ang2), a regulator of angiogenesis, is stored with other pro-inflammatory and pro-thrombotic mediators, in endothelial-specific vesicles called Weibel-Palade bodies (WPBs). The WPB secretagogue, histamine, delays Ang2 secretion by activating Rab46-specific trafficking of Ang2-positive WPBs to the microtubule organising centre (MTOC), where they persist until Ca^2+^ binds to the EF-hand of Rab46, enabling detachment. Here, using Ca^2+^ imaging and high-resolution light microscopy, we pharmacologically investigated the contribution of endolysosomal two-pore channels proteins (TPC) to the Ca^2+^ signal necessary for Ang2 secretion. We show an increase in the histamine-evoked clustering of Rab46 (and thus WPBs) at the MTOC in the presence of TPC inhibitors Ned19 and tetrandrine, and a decrease in the presence of a TPC2 agonist, TPC2-A1-N. Histamine-evoked secretion of Ang2 was decreased by pharmacological inhibition of TPC channels but potentiated in the presence of an TPC2 agonist. These data suggest that histamine-mediated Ca^2+^ release via TPC2 channels is necessary for the Rab46-dependent detachment of Ang2-positive WPBs from the MTOC and thus Ang2 secretion.

**Summary:** Ca^2+^ binding to the EF-hand of Rab46 in endothelial cells has previously been reported but the molecular mechanisms and functional relevance is unclear. Here the authors show that Ca^2+^ released from TPC2 regulates the detachment of Rab46 from the MTOC and thereby allows secretion of Ang2 from WPBs.

## Introduction

Endothelial cells (ECs) play a crucial role in the maintenance of vascular health by dynamically modulating vascular tone, angiogenesis, inflammation and haemostasis [1]. ECs store a plethora of proinflammatory and prothrombotic cargo in EC-specific organelles termed Weibel-Palade bodies (WPBs) [2] [3] [4]. During vascular injury, WPBs undergo rapid exocytosis and secrete their cargo into the vascular lumen to instigate vascular repair. For example: ejection of von Willebrand factor (vWF) initiates platelet plug formation [5]; cell surface expression of P-selectin attracts leukocytes [6] and release of Ang2 induces EC migration [7]. However, variation in the recruitment and secretion of cargo confers a plasticity on WPBs that enables them to impart functionally appropriate responses to non-injury stimuli [8] [9] [10]. This plasticity is vital so that processes such as inflammation and angiogenesis can be controlled without the risk of thrombosis. We have previously described a unique stimuli-dependent trafficking pathway that regulates WPB cargo secretion [8]. Acute histamine (immunogenic), but not thrombin (injury), signalling evokes trafficking of a subpopulation of WPBs away from the plasma membrane to the microtubule organising centre (MTOC). This subset of WPBs carries Ang2, a proangiogenic protein that regulates EC migration and vascular remodelling. The histamine-evoked localisation of WPBs at the MTOC limits Ang2 exocytosis (as compared to a thrombin response), whilst, WPBs storing pro-inflammatory signals, such as P-selectin are trafficked to the plasma membrane and their cargo secreted. This pathway provides a histamine appropriate immune response by regulating the selective secretion of WPB cargo thus preventing superfluous angiogenesis. This unique trafficking pathway is dependent on a Rab GTPase, Rab46 [8] [11] [12].

The Rab family of small GTPases are master regulators of membrane trafficking [13]. Rab46 is an unusual Rab GTPase because in addition to a highly conserved Rab domain, it also has Ca^2+^-sensing (EF-hand) domains [11] [8]. Rab46 is one of two functional proteins translated from the *EFCAB4B* gene; the other isoform being a non-Rab Ca^2+^ sensor involved in store-operated Ca^2+^ entry in T-cells (CRACR2A) [14]. In human umbilical vein endothelial cells (HUVECs), Rab46 is necessary for the histamine-evoked trafficking of Ang2-positive WPBs to the MTOC [8]. This Rab46-regulated retrograde trafficking is dependent on microtubules and dynein; indeed, Rab46 has been shown to be a direct dynein adaptor in endothelial and T cells [12, 15]. However, in ECs, as opposed to T-cells, the Rab46 and dynein mediated retrograde trafficking towards the MTOC in response to histamine is independent of Ca^2+^ [8, 12]. Moreover, Ca^2+^ binding to the EF-hand of Rab46 is necessary for detachment and dispersal of WPBs away from the MTOC. We therefore propose that histamine-evoked localised Ca^2+^ signals regulate Ang2 secretion by liberating Ang2-positive WPBs clustered at the MTOC. Understanding the mechanisms underlying secretion is important because inappropriate Ang2 serum levels promote cardiovascular diseases [16] [17], including myocardial infarction [18] and heart failure [19]. Furthermore, studies suggest that Ang2 is associated with a greater risk of cardiovascular mortality [20] and is a predictor of mortality in myocardial infarction patients [21].

Many WPB secretagogues, including histamine and thrombin, mobilise Ca^2+^ from intracellular stores [22] [23] [24]. The best-characterized intracellular Ca^2+^ pathway is that mediated by IP_3_ produced by receptor-evoked activation of phospholipase C. IP_3_ targets Ca^2+^ channels (IP_3_ receptors) present on the endoplasmic reticulum (ER) [25]. Indeed, both histamine and thrombin evoke release of Ca^2+^ from ER stores via IP_3_ and the subsequent rise in intracellular Ca^2+^ is necessary and sufficient for WPB cargo secretion. We therefore questioned the unique nature of the Ca^2+^ signal evoked by histamine that is necessary for Rab46-dependent detachment of WPBs from the MTOC, presuming it must be distinct to Ca^2+^ released from the ER. Highly localised Ca^2+^ nanodomain signalling is a characteristic of the second messenger Nicotinic Acid Adenine Dinucleotide Phosphate (NAADP) [26]. Major research effort in the past two decades has shown that the mode of action of NAADP differs from the canonical IP_3_-ER release axis. Generated in response to various receptor-agonists, NAADP triggers Ca^2+^ release from thapsigargin-insensitive stores that have been identified as the endo-lysosomal Ca^2+^ system [27]. Due to the varied and dynamic localisation of the endo-lysosomes around the cell, the Ca^2+^ released is small but highly localised. In some cell types this NAADP-dependent Ca^2+^ release on its own is sufficient to mediate a cellular function such as phagocytosis in macrophages [26] or stimulus-coupled secretion in T cells [28]. However, in pancreatic acinar cells [29], the Ca^2+^ released from NAADP-sensitive stores serves as a co-agonist, or a trigger, to the ER localised IP_3_R (also known as ‘the trigger hypothesis’) thereby modulating the global Ca^2+^ response by Ca^2+^-induced Ca^2+^ release (CICR) [30]. In ECs several agonists are coupled to NAADP: Esposito *et al* showed that in ECs histamine H_1_R is coupled to NAADP and inhibition of NAADP signalling decreases vWF release evoked by H_1_R agonism [31]. We hypothesized that upon histamine signalling, NAADP-dependent release of Ca^2+^ is necessary for regulating detachment of Rab46 (and thus WPBs) from the MTOC.

Here, we explored the relationship between Ca^2+^ released from NAADP sensitive two pore channel proteins 1 and 2 (TPC1/2) and Rab46-dependent trafficking using pharmacological antagonists (Ned19 and tetrandrine), a novel cell-permeable activator of TPC2 (TPC2-A1-N) and high-resolution microscopy. We demonstrate for the first time that a Rab GTPase is a context-dependent sensor of Ca^2+^ released from the endolysosomal cation channel TPC2 and this has important roles in regulating Ang2 secretion, a key regulator of angiogenesis.

## Results

### Ned19 promotes histamine-evoked Rab46 retention at the MTOC

Ca^2+^ binding to the EF-hand of Rab46 is necessary for WPB detachment from the MTOC [8]. Thereby, anticipating that this signal must be distinct from Ca^2+^ released from the ER, we investigated the impact of pharmacological modulation of the NAADP-sensitive TPC1/2 channels, on Rab46 (and thus WPB) retention at the MTOC. Here, we used human Aortic Endothelial Cells (hAECs) as a physiologically relevant EC type. Therefore (due to cell type disparity), we first confirmed that histamine-evoked retrograde clustering of Rab46 at the MTOC was consistent with HUVECs and was independent of Ca^2+^ (Fig. 1 and Supplementary Fig. 1).

**Figure 1.**
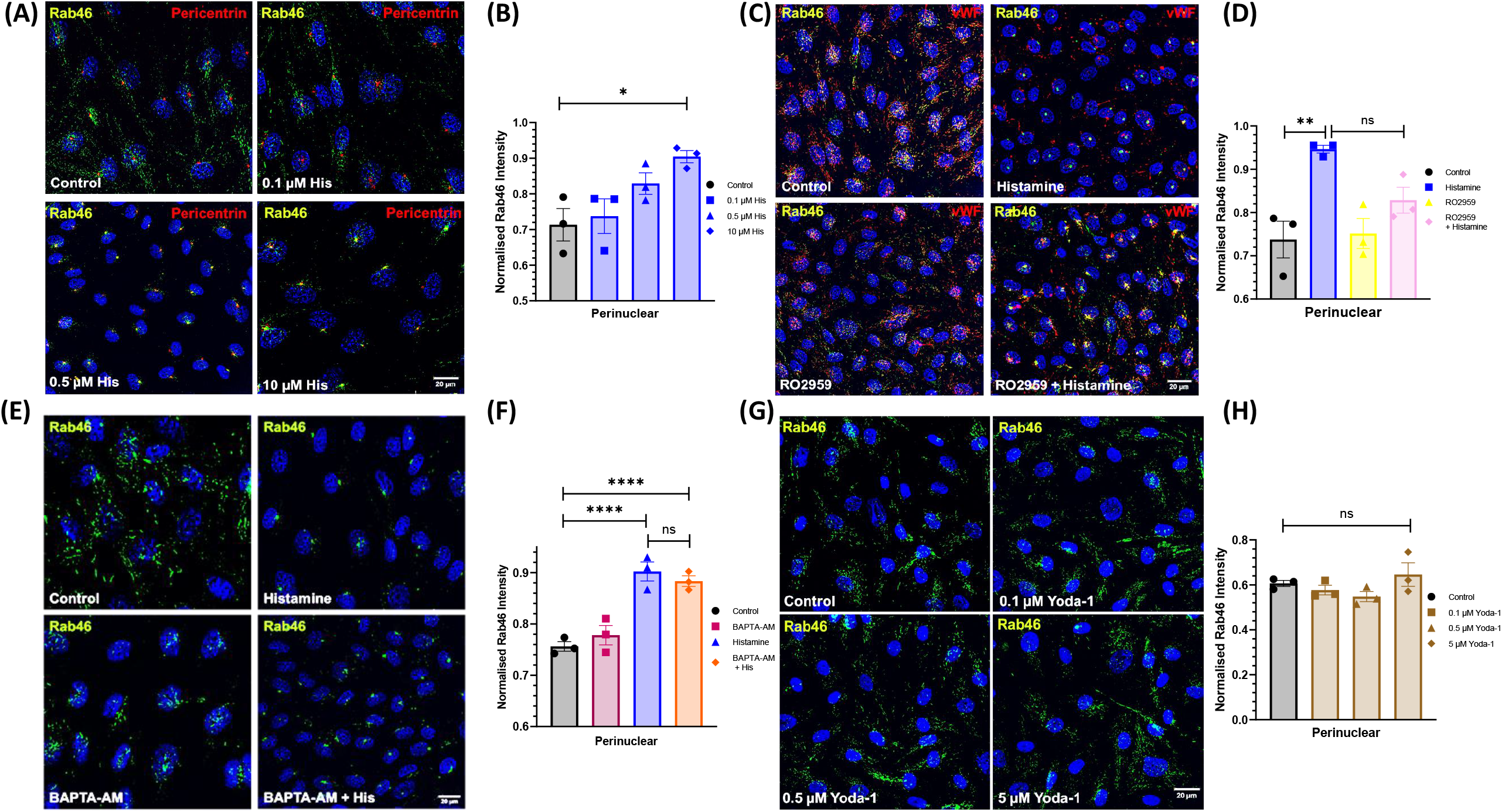
Histamine-evoked trafficking of WPBs to the MTOC is independent of Ca^2+^ in human aortic endothelial cells. (A, B) Representative images (A) and mean perinuclear distribution (0 – 2 μM from the nucleus) (B) of hAECs immunostained for Rab46 (green) and pericentrin (red) as a marker of the MTOC in response to 10 mins treatment of increasing concentrations of histamine. Blue = Hoechst (nuclei). Scale bar = 20 μM and applies to all images. (C, D) Representative images (C) and mean perinuclear distribution of Rab46 (D) of hAECs immunostained for Rab46 (green) and vWF (red) as a marker of WPBs in response to 10 mins treatment with 30 μM histamine in hAECs preincubated with vehicle control or 10 μM of the Orai1 channel inhibitor RO5929. (E, F) Representative images (E) and mean perinuclear distribution (F) of hAECs immunostained for Rab46 (green) in response to 10 mins treatment with 30 μM histamine in hAECs preincubated with vehicle control or 20 μM of the Ca^2+^ chelator BAPTA-AM. (G, H) Representative images (G) and mean perinuclear distribution (H) of hAECs immunostained for Rab46 (green) in response to 10 mins treatment with increasing concentrations of the Piezo1 Ca^2+^ channel activator Yoda1. All distribution plots represent mean ±S.E.M normalised intensity. Each dot represents the mean of an independent experiment (n/N=3/18-30). * p<0.05, ** p<0.01, *** = p<0.001, **** = <0.0001 from One-way ANOVA and Tukey post-hoc analysis.

Next, to determine whether NAADP-sensitive channels are the source of Ca^2+^ required for detachment from the MTOC, we first explored the Ca^2+^ signal evoked by histamine in hAECs. The histamine concentration-dependent Ca^2+^ curve displayed a rightward shift when hAECs were pretreated with the NAADP channel antagonist *Trans*-Ned19 (Ned19: [32]) (Fig. 2A) indicating that histamine evokes Ca^2+^ release from TPC1 or TPC2 (NAADP-sensitive channels). At a low concentration of histamine (0.1 μM), the Ca^2+^ rise is prevented by Ned19 (Fig. 2B, C, D. Supplementary Fig. 2A, B), suggesting that low concentrations, of histamine trigger Ca^2+^ mobilisation solely from NAADP-sensitive intracellular Ca^2+^ stores. Conversely, at 30 μM histamine NAADP antagonism only decreased the peak amplitude of the histamine-evoked Ca^2+^ release, without completely inhibiting it (Fig. 2E, F, G), suggesting that the peak Ca^2+^ response at higher concentrations of histamine is a combination of Ca^2+^ from both NAADP and other (potentially IP_3_-sensitive) stores. Yoda1 activation of the Piezo1 Ca^2+^ channel induces a global intracellular Ca^2+^signal that was not affected by Ned19 (Fig. 2H).

**Figure 2.**
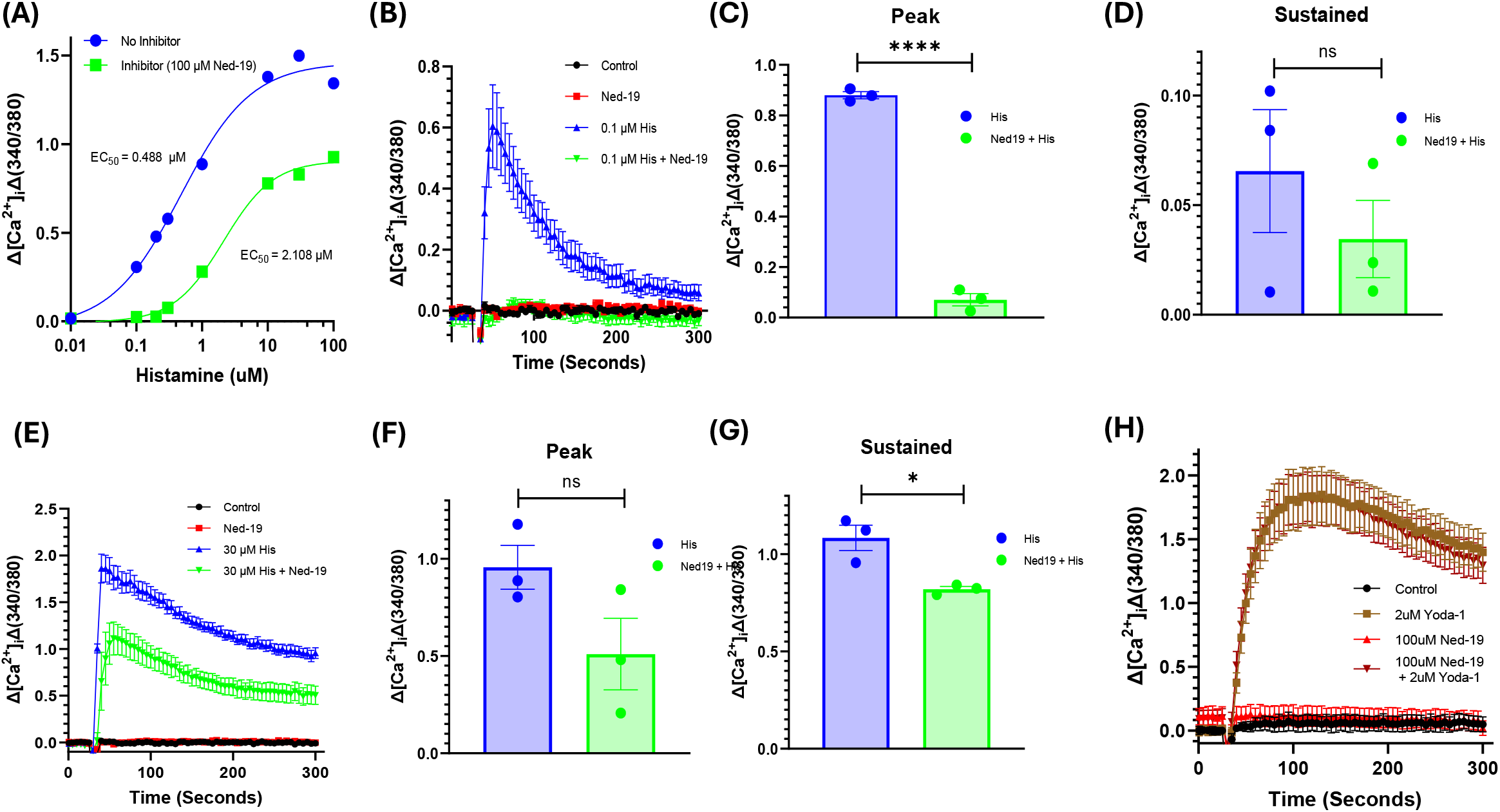
TPC1/2 antagonism inhibits histamine-evoked Ca^2+^ mobilisation in hAECs. (A) hAECs demonstrate a rightward shift in the histamine-evoked concentration curve when preincubated with 100 μM Ned19. (B) Representative traces of changes in Ca^2+^ concentration in hAECs evoked by application of 0.1 µM histamine at 60 seconds where cells were pre-treated 30 mins with 100 µM Ned19 or vehicle control. Recordings were obtained in 1.5 mM Ca^2+^ buffer unless otherwise stated. (C) Mean peak amplitude and (D) mean sustained responses in hAECs of the traces obtained in (B). (E) Representative traces of Ca^2+^ mobilisation in hAECs evoked by application of 30 µM histamine at 60 seconds where cells were pre-treated 30 mins with 100 µM Ned19 or vehicle control. (F) Mean peak amplitude and (G) mean sustained responses in hAECs of the Ca^2+^ traces obtained in (E). (H) Representative traces of Ca^2+^ release in hAECs evoked by application of 2 µM Yoda1 at 60 seconds where cells were pre-treated with 100 µM Ned19 or control for 30 minutes. n/N=3/9, error bars represent SEM, *** p value < 0.001, **** p value < 0.0001 by One-way ANOVA with Tukey post-hoc analysis or unpaired t-test.

**Figure 3.**
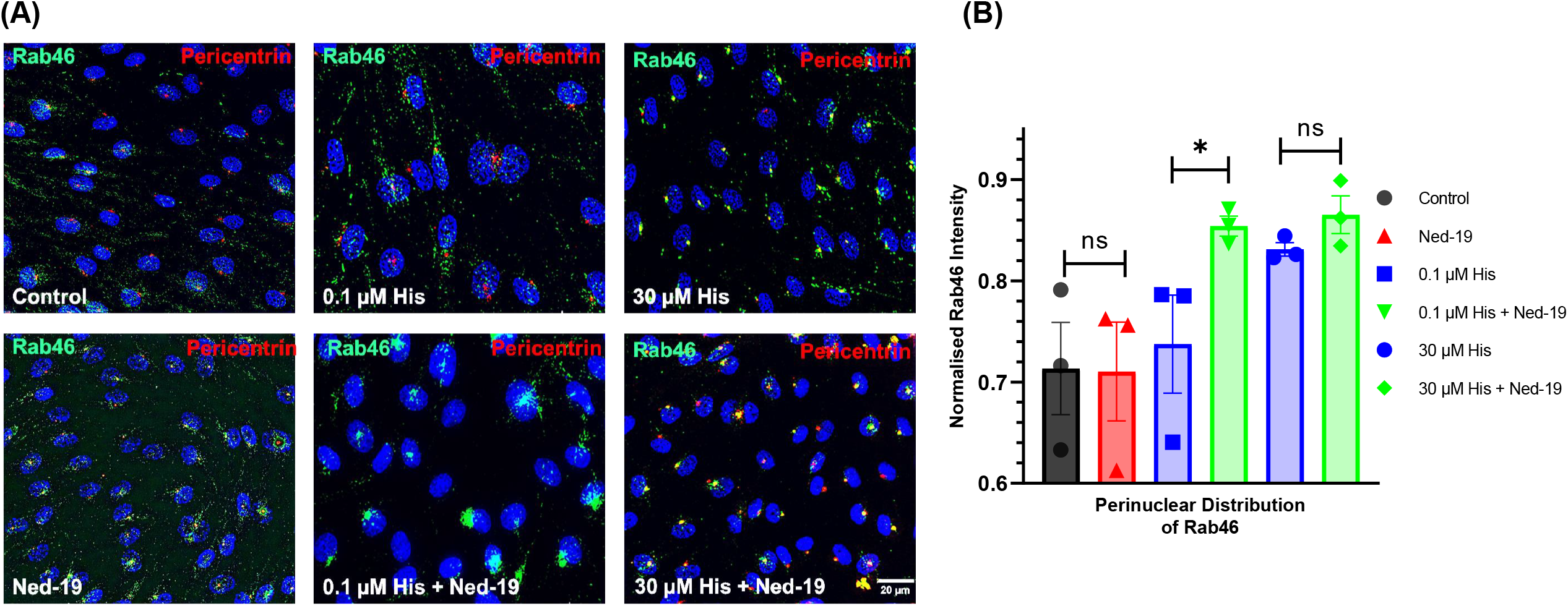
Inhibition of Ca^2+^ release from TPC1/2 promotes histamine evoked Rab46 clustering at the MTOC. (A) Representative images of hAECs immunostained for Rab46 (green) and pericentrin (red) stimulated with 0.1 μM or 30 μM histamine (or vehicle control) in the presence or absence of 100 μM Ned19. Scale bar = 20 μm applies to all images. Blue = Hoechst (nuclei). (B) Quantitative analysis as described in the methods of Rab46 perinuclear distribution in hAECs depicted in images (A). n/N=3/15. *p-value < 0.05 from One-way ANOVA with Tukey post-hoc analysis.

We hypothesized that histamine-evoked localised Ca^2+^ release from NAADP-sensitive TPC1 or TPC2 is necessary and sufficient for the detachment of Rab46 from the MTOC. Thereby, inhibition of TPC1/2 should enhance Rab46 clustering at this locale. Endogenous Rab46 localisation was observed, and fluorescence intensity quantified in hAECs pretreated with Ned19 (or control) prior to histamine stimulation. Ned19 inhibition of TPC1/2-mediated Ca^2+^ release enhanced the clustering of Rab46 at the MTOC in the presence of low histamine concentration (Fig. A, B. Supplementary Fig. 3). This data indicates that Ca^2+^ released from TPC1 or TPC2 channels is necessary for Rab46 detachment from the MTOC.

### Inhibition of TPC2 potentiates histamine-evoked clusters of Rab46 at the MTOC

To distinguish the contribution of TPC1 or TPC2 channels to Rab46 trafficking, we investigated the effects of pre-incubating cells with the Ca^2+^ channel blocking drug tetrandrine. Tetrandrine has been shown to antagonize both NAADP- and PI(3,5)P_2_-mediated signalling at TPC2 [33]. Pre-incubation of hAECs with tetrandrine inhibited the histamine induced Ca^2+^ response (Fig. 4A - F) and increased the clustering of Rab46 at the MTOC (Fig. 4G, H). These data suggest that histamine evokes Ca^2+^ release from the NAADP-sensitive channel TPC2 and this regulates the Ca^2+^-dependent detachment of Rab46 from the MTOC. To further support this hypothesis, we explored the effects of a TPC2-specific agonist [34].

**Figure 4.**
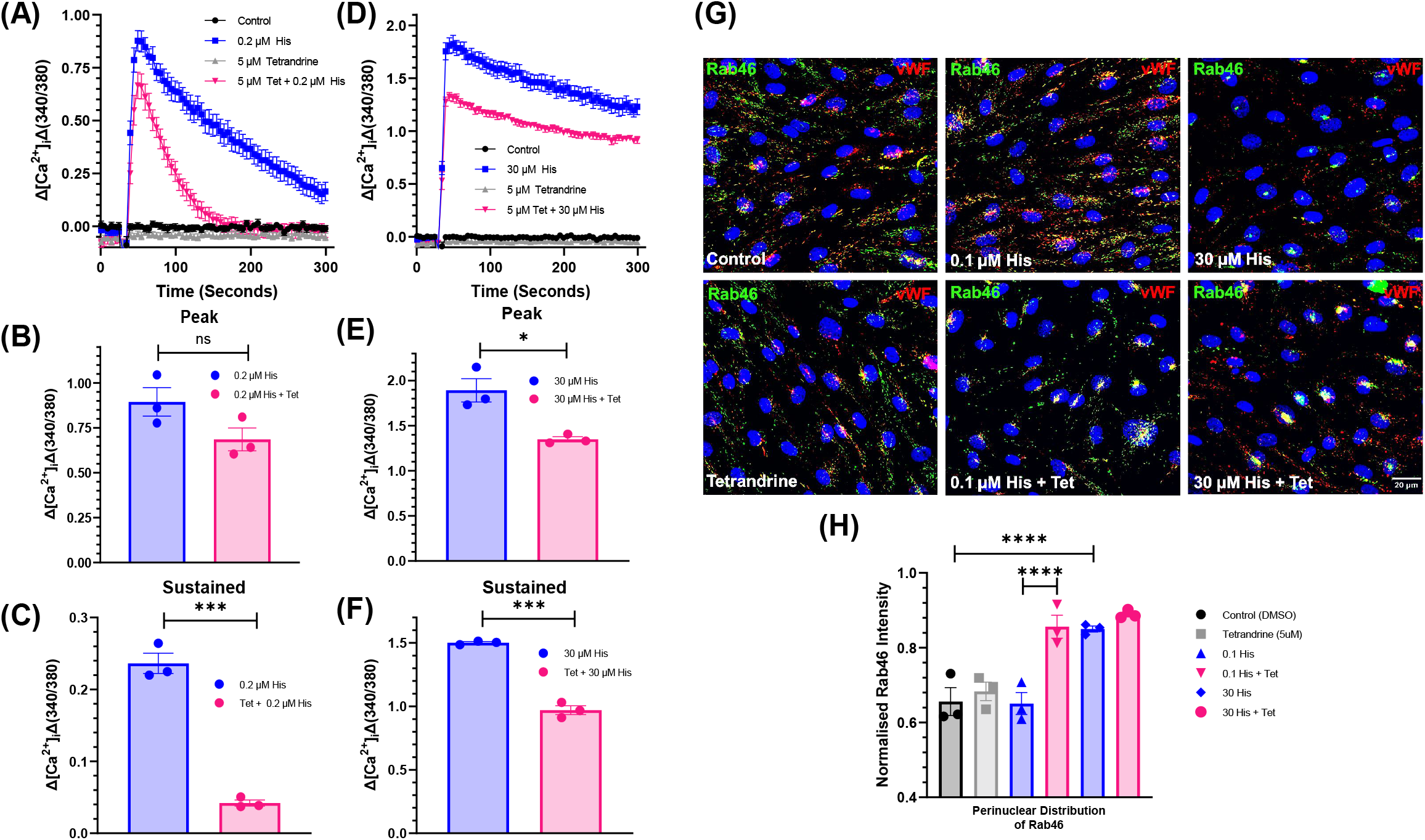
The TPC2 antagonist tetrandrine potentiates histamine-evoked Rab46 clustering at the MTOC. A) Representative traces of Ca^2+^ mobilisation in hAECs evoked by application of 0.2 µM histamine at 60 seconds where cells were pre-treated 30 mins with 5 µM tetrandrine or vehicle control. (B) Mean peak amplitude and (C) mean sustained responses in hAECs of the Ca^2+^ traces obtained in (A). (D) Representative traces of Ca^2+^ mobilisation in hAECs evoked by application of 30 µM histamine at 60 seconds where cells were pre-treated 30 mins with 5 µM tetrandrine or vehicle control. (E) Mean peak amplitude and (F) mean sustained responses in hAECs of the Ca^2+^ traces obtained in (D). n/N = 3/9. (G) Representative images of hAECs immunostained for Rab46 (green) and pericentrin (red) as a marker of the MTOC in response to 10 mins treatment of 0.1 or 30 μM histamine in the absence or presence of 5 μM tetrandrine. Blue = Hoechst. Scale bar = 20 μM applies to all images. (H) Mean data of the perinuclear localisation of Rab46 in response to 10 mins stimulation with histamine in cells preincubated with vehicle control or 5 μM tetrandrine. n/N = 3/15. *p-value < 0.05 from One-way ANOVA and unpaired t-tests.

### A TPC2 agonist prevents histamine-evoked clusters

The selective cell-permeable TPC2 agonists TPC2-A1-N and TPC2-A1-P mimic NAADP (Ca^2+^ selective) and PI(3, 5)P_2_ (Na^+^ selective) action respectively [35]. TPC2-A1-N, but not TPC2-A1-P, (Fig. 5A. Supplementary Fig. 5A, B, C) evoked a concentration-dependent rise of Ca^2+^ in hAECs and HUVECs (Supplementary Fig. 5A). The biasing of TPC2 to a Ca^2+^ permeable channel by TPC2-A1-N sensitises local Ca^2+^ signals to IP_3_ [36]. Therefore, we questioned if TPC2-A1-N could potentiate the localised Ca^2+^ signal evoked by histamine and if an enhanced Ca^2+^ signal at the MTOC could provoke detachment of Rab46 WPBs from the MTOC. Pre-treatment of hAECs with 30 μM TPC2-A1-N inhibits the histamine-evoked response (Fig. 5B) indicating depletion of ER stores (as shown previously the Ca^2+^ signal derived from high concentrations of TPC2-A1-N triggers CICR [37]). To determine if a lower concentration (10 μM) of TPC2-A1-N could evoke a localised Ca^2+^ signal (which, when used alone in this timeframe, is not detectable by Fura2 epifluorescence measurements (Fig. 5A)), we considered context-dependence (i.e. are physiological stimuli necessary for enabling the close association between TPCs and the Ca^2+^ sensing proteins necessary for downstream events). ECs stimulated with 10 μM TPC2-A1-N after activation with histamine exhibited a further Ca^2+^ peak (Fig. 5C) suggesting 10 μM TPC2-A1-N activates a localised response. Stimulation of hAECs with 10 μM TPC2-A1-N in the presence of histamine potentiates the sustained Ca^2+^ signal as compared to histamine alone (Fig. 5D -F). To determine if this enhanced Ca^2+^ signal promotes Rab46 dispersal from the MTOC, we treated hAECs with histamine in the presence and absence of TPC2-A1-N and quantified Rab46 cellular distribution. TPC2-A1-N stimulation alone did not evoke trafficking of Rab46, or decrease the Rab46 signal, but inhibited the histamine-evoked clustering of Rab46 at the MTOC compared to histamine alone (Fig. 5G, H, Supplementary Fig. 5D). This further indicates that Ca^2+^ release from TPC2 promotes the detachment of Rab46 from the MTOC.

**Figure 5.**
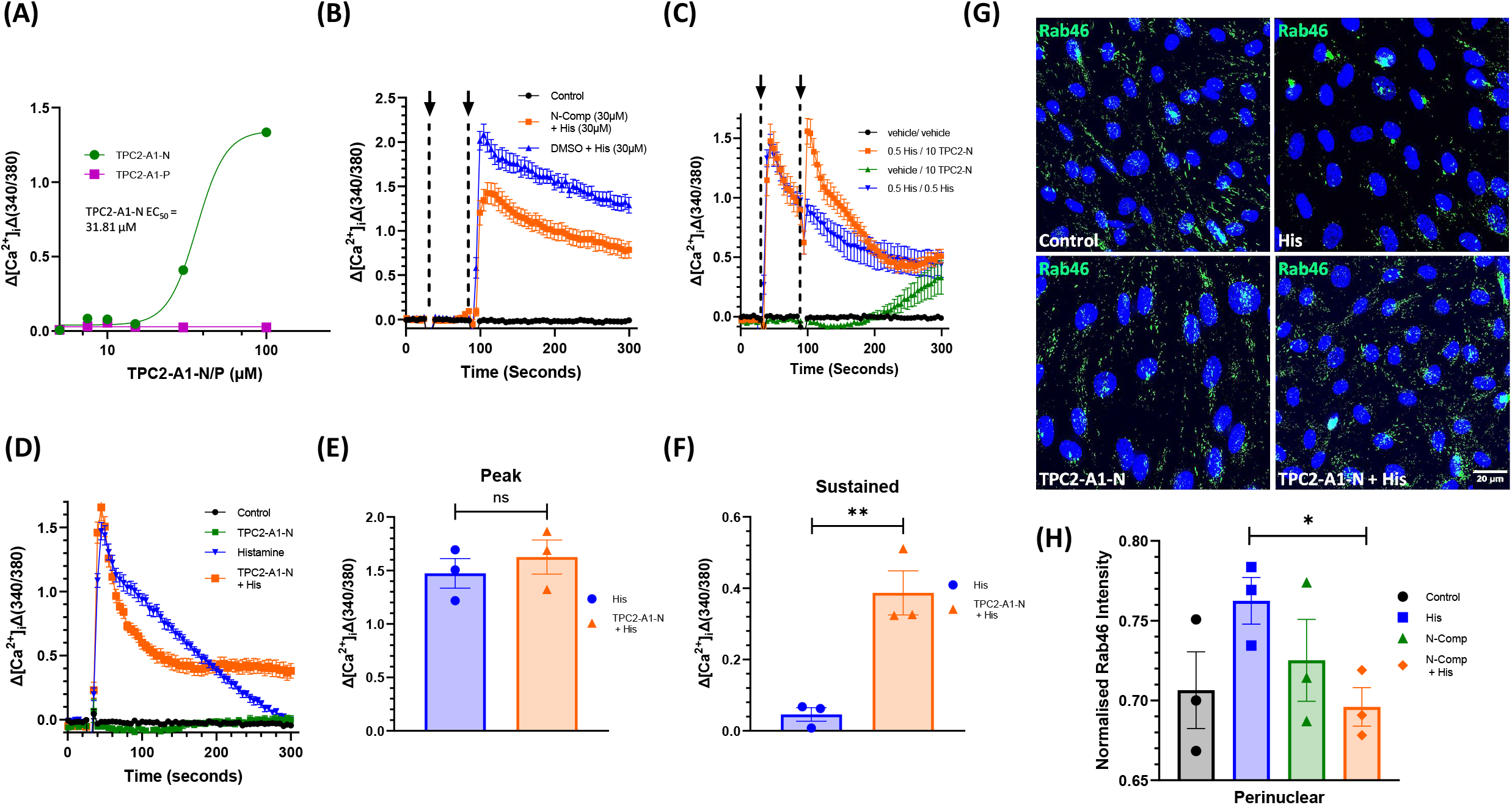
TPC2 activation enhances Rab46 trafficking. (A) Mean data of hAECs treated with (0.1-100 µM) TPC2-A1-N or TPC2-A1-P displayed as concentration-response curves. n/N = 3/9. (B) Representative traces measuring changes (Δ) in intracellular Ca^2+^ evoked by 30 μM histamine (arrow at 60 secs) after pre-stimulation with 30 μM TPC2-A1-N or control (10 secs). n/N = 3/9. (C) Representative traces measuring changes (Δ) in intracellular Ca^2+^ evoked by 10 μM TPC2-A1-N (arrow at 60 secs) after pre-stimulation with 0.5 μM histamine or control (10 secs) n/N = 3/9. (D-F). Representative traces (D) and mean data (E, F) measuring changes (Δ) in intracellular Ca^2+^ evoked by 0.5 μM histamine or simultaneous addition of 0.5 μM histamine and 10 μM TPC2-A1-N. Recorded in 0 Ca^2+^buffer. n/N = 3/9. (G, H) Representative images (G) and mean distribution analysis (H) of Rab46 (green) in hAECs treated with 0.5 μM histamine alone or simultaneous addition of 0.5 μM histamine and 10 μM TPC2-A1-N. Simultaneous addition decreased Rab46 (green) clusters at the MTOC. Scale bar = 20 μM. Blue = Hoechst (nuclei). n/N=3/15, error bars represent SEM, *** p value < 0.001, **** p value < 0.0001 by One-way ANOVA with Tukey post-hoc analysis or unpaired t-test.

### TPC2 regulated Rab46 trafficking regulates histamine evokes Ang2 secretion from WPBs

The subset of WPBs clustered at the MTOC in response to histamine are Ang2-positive [8]. Thereby, to probe the functional relevance of Ca^2-^ dependent detachment of WPBs from the MTOC, we examined the role of TPC1/2 Ca^2+^ release on histamine-evoked Ang2 secretion. First, we show that mimicking vascular injury and exposing hAECs to a combination of thrombin and histamine (combo) evokes Ang2 secretion (Fig. 6A). Consistent with our previous analysis, Ang2 secretion is not augmented by higher concentrations of histamine (Fig. 6A). Yoda1 mediated Ca^2+^ influx does not evoke Ang2 secretion. Inhibition of NAADP-mediated Ca^2+^ release by Ned19, inhibits histamine-evoked Ang2 secretion, to basal levels (Fig. 6A). We questioned if potentiation of TPC2 dependent Ca^2+^ release (as shown in Fig. 5D) and the subsequent dispersal of Rab46 from the MTOC (Fig. 5H) could promote the secretion of Ang2 upon histamine signalling. Whilst neither 10 nor 30 μM TPC2-A1-N alone evoked secretion of Ang2 (Fig. 6B), the stimulation of hAECs with TPC2-A1-N in the presence of histamine (0.5 His + 10N) potentiated secretion of Ang2 compared to histamine (0.5 His) alone. These data suggest that histamine evokes trafficking of WPBs to the MTOC where the Ca^2+^ released from TPC2 elicits detachment of Ang2-positive from the microtubules, thus allowing secretion of Ang2.

**Figure 6.**
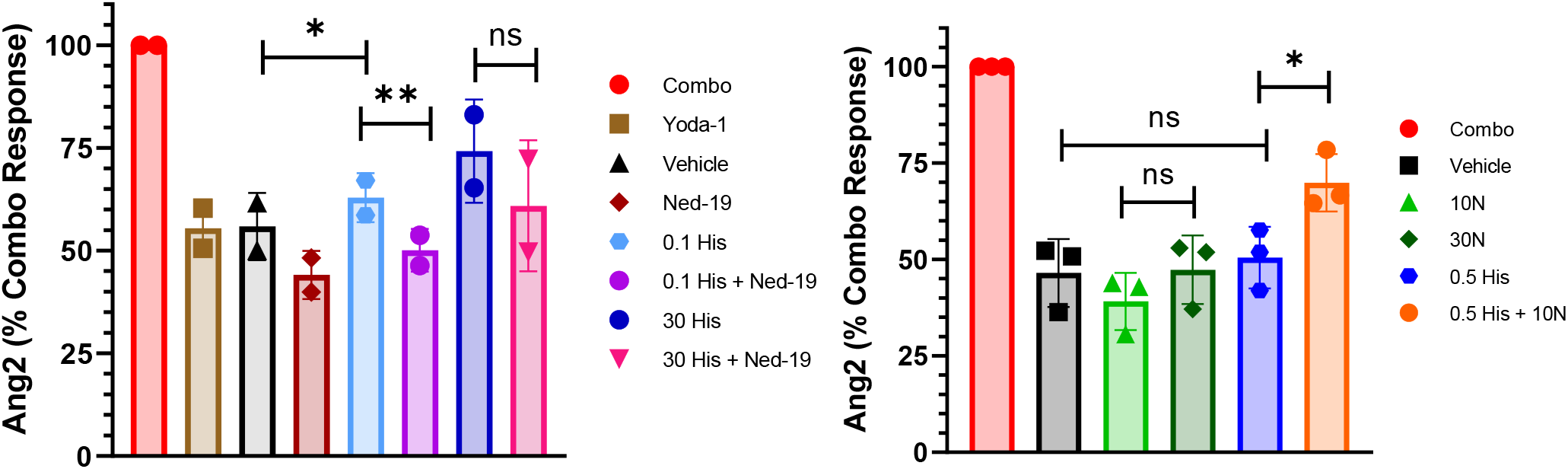
TPC2-dependent detachment of WPBs regulates histamine-evoked Ang2 secretion in ECs. (A) ELISA-based analysis of protein levels for Ang2 in the media collected from hAECs following exposure to histamine (0.1 μM or 30 μM His) alone or histamine following incubation with 100 μM Ned19 (0.1 His + Ned-19, or 30 His + Ned-19). In addition, hAECs were exposed to the plasma membrane cation channel agonist, Yoda1 (2 μM) to indicate specificity for TPC1/2 activity. Results expressed as a percentage of Ang2 levels measured in hAECs exposed to a combination of 2.5 U thrombin and 30 μM histamine (combo: positive control). n/N 2/6. (B) ELISA-based analysis of protein levels for Ang2 in the media collected from hAECs following exposure to 0.5 μM histamine or TPC2-A1-N (10 μM N, 30 μM N) alone or histamine and TPC2-A1-N simultaneously (0.5 His + 10N). Results expressed as a percentage of Ang2 levels measured in the media collected from hAECs exposed to a combination of 2.5 U thrombin and 30 μM histamine (combo: positive control). n/N 3/9. *p-value < 0.05 from 2-way ANOVA with Tukey post-hoc analysis.

## Discussion

Rab46 is a Ca^2+^-sensing Rab GTPase localised to a subpopulation of WPBs in endothelial cells. We have previously demonstrated in HUVECs that acute histamine signalling evokes a Rab46 and dynein-dependent trafficking of Ang2-positive WPBs to the MTOC and that once localised to the MTOC, Ca^2+^ binding to the EF-hand of Rab46 is necessary for WPB to detach [8]. Using a pharmacological approach, this work describes how, in hAECs, TPC2 provides the localised Ca^2+^ signal necessary for the release of Rab46 from the MTOC and this has functional relevance upon physiological signalling. Inhibition of TPC1/2-evoked Ca^2+^ release by both tetrandrine and Ned19 results in enhanced histamine-evoked clustering of Rab46 at the MTOC, suggesting the availability of Ca^2+^ is reduced at this locale. Increased clustering of Rab46 at the MTOC inhibits the secretion of Ang2 from WPBs. Accordingly, enhanced TPC2-mediated Ca^2+^ release promotes detachment of Rab46 from the MTOC in the presence of histamine, which is functionally relevant because not only does it allow dispersal of WPBs but, potentiates secretion of Ang2.

Considering the need for rapid exocytosis and secretion of WPB cargo in response to vascular injury, we had originally surmised that an EF-hand-containing Rab GTPase localised to WPBs would expediate this Ca^2+^-dependent process, thus overriding the requirement for intracellular signalling events. However, in HUVECs, Rab46 was not activated upon thrombin (injury) signals, but was necessary for dynein-dependent trafficking of a subpopulation of WPBs containing Ang2 to the MTOC upon acute histamine stimulation, thus limiting Ang2 secretion [8]. In agreement, here histamine evokes concentration-dependent trafficking of Rab46 to the MTOC in hAECs. We demonstrated that Rab46 clustering is localised to pericentrin, a marker for the MTOC, and our automated (Supplementary Appendix 1) cellular distribution analysis confirms a histamine-dependent movement of Rab46 fluorescent intensity from the intermediary region of the cell (2-5 μM from the nucleus) to the perinuclear (0-2 μM from the nucleus). Whilst Wang *et al* proposed that in T-cells Rab46 recruitment of dynein was dependent upon Ca^2+^ [15], our studies support a Ca^2+^-independent mechanism in both HUVECs using EF-hand mutants [8] [12] and here in hAECs where inhibition of Ca^2+^ influx through the store-operated Orai1 channel or chelation of intracellular Ca^2+^ did not inhibit histamine-evoked retrograde trafficking. Moreover, promoting a global rise in intracellular Ca^2+^ by stimulating the plasma membrane cation channel, Piezo1, does not induce trafficking of WPBs. However, we have shown that Ca^2+^ is necessary for the detachment of WPBs from the MTOC and this requires binding to the EF-hand of Rab46 [8]. Since both histamine and thrombin evoke global intracellular Ca^2+^ signals by stimulating the release of Ca^2+^ from ER stores with subsequent influx through the Orai1 channel (SOCE), we questioned the molecular mechanisms underlying the Ca^2+^ release that is restricted to a nanodomain around the MTOC and is specific to histamine signalling.

In addition to ER stores, the endolysosomal acidic stores are known to modulate intracellular free Ca^2+^. NAADP is a second messenger that acts on two pore channels (TPCs) indirectly by binding to intermediary accessory proteins to induce release of Ca^2+^ from endolysosomal stores [38]. Two channel isoforms are expressed in humans, TPC1 and TPC2 [39] and they have been implicated in the generation of NAADP-mediated Ca^2+^ signals in endothelial cells [31] [40]. Ca^2+^ released from these channels can trigger Ca^2+^-induced Ca^2+^ release from the ER which will evoke a global Ca^2+^signal in the cytosol. However, it is evident that there is a localised signal around the channels themselves owing to restricted Ca^2+^ diffusion. This patterning of free Ca^2+^ enables the formation of discrete nanodomains where Ca^2+^ and Ca^2+^ binding proteins are uniquely positioned, thus enabling coupling of stimuli to appropriate Ca^2+^-dependent cellular responses such as phagocytosis in macrophages [26]. Due to the restricted positioning of Rab46 and WPBs at the MTOC upon histamine stimulation and the need for histamine to be distinguished from other Ca^2+^ mobilising stimuli, we had questioned if TPC1/2 provided the Ca^2+^ necessary for detachment. The TPC1/2 antagonist Ned19 inhibited histamine-evoked Ca^2+^ signalling but not the global Ca^2+^ rise promoted by activation of a plasma membrane cation channel. Ned19-mediated inhibition of Ca^2+^ release was sufficient to cause an increase in the retention of Rab46 at the MTOC upon histamine stimulation, which inhibited Ang2 secretion. The data suggests that the source of free Ca^2+^ that is necessary for detachment of Rab46, and thus WPBs, from the MTOC, is from TPC1/2 activity. Whilst Ned19 has previously been shown to inhibit histamine and VEGF (but not thrombin) -evoked Ca^2+^ signals in ECs [31] [40] (which ultimately impacted secretion of vWF and angiogenesis), the molecular details of this circuitry are lacking.

Another functional distinction of NAADP Ca^2+^ signalling is that it exhibits a bell-shaped concentration response relationship; meaning that nM concentrations of intracellular NAADP, evokes Ca^2+^ release but release is not detectable at μM concentrations [41]. Studies in Jurkat T-cell lines showed maximal Ca^2+^ release upon stimulation with 100 nM NAADP whilst concentrations of >1 μM failed to elicit any response whilst inhibiting T-cell receptor activation [42]. Analysis in different cell types, including pulmonary smooth muscle cells [43] and ventricular cardiac myocytes [44] have verified this phenomenon. In this study and in HUVECs [8] we observe maximal redistribution of Rab46 to the MTOC (perinuclear region in our distribution analysis) with histamine concentrations > 10 μM. We postulate that histamine either generates NAADP at self-inhibitory levels or, as our data suggests, the lysosomal stores have become deplete of Ca^2+^ at concentrations > 10 μM and thus the effects of TPC1/2 antagonism are more effective at lower doses.

TPC2 has previously been shown to be predominantly localised to acidic late-endosomes/lysosomes (typically localised around the MTOC), whilst TPC1 is found on recycling endosomes and early endosomes in the cytosol [45, 46], suggesting a prominent role for TPC2 in this pathway. Tetrandrine is a broad-spectrum channel blocker used to inhibit TPC2 [35, 47] and to a lesser extent TPC1 [48]. With its potency towards TPC2, it is used here to discriminate between these channels. Tetrandrine inhibited the histamine-evoked Ca^2+^ response and evoked increased clustering of Rab46, indicating that TPC2 provides the free Ca^2+^ necessary for WPB detachment. Future studies using super-resolution imaging to exactly determine the spatial resolution of TPCs and Rab46 would be valuable, with the caveat that there are presently no suitable TPC channel antibodies and overexpression of GFP-tagged channels may impact the balance between Ca^2+^ availability and trafficking events. In addition, the use of plant flavonoids such as pratensein and duartin, which block TPC2 [49], should be used to verify the role of TPC2.

The recent discovery of stable, cell-permeant agonists that are selective for TPC2 over TPC1 and TRPML allowed further determination of the role of TPC2 [35]. The agonists mimic either NAADP (TPC2-A1-<u>N</u>) or PI(3,5)P_2_ (TPC2-A1-<u>P</u>) binding and confer a selectivity on the channel to either Ca^2+^ or Na^+^ respectively. Yuan *et al* indicated that the action of TPC2-A1-N is distinct to NAADP and its binding proteins, is not inhibited by Ned19 and does not display a bell-shaped dose response curve. Moreover, this TPC2 agonist sensitises cells to IP_3_ [37]. Therefore, we postulated that TPC2-A1-N could potentiate the Ca^2+^ response of histamine, a physiological activator of IP_3_. Since, the majority of the signal evoked by TPC2-A1-N involves Ca^2+^ release from the ER, we used a relatively low concentration of TPC2-A1-N (10 μM) where there was no detectable effect on cytosolic global Ca^2+^ concentration over the time frame of acute histamine stimulation. However, the efficacy of 10 μM was observed when used in the presence of submaximal histamine stimulation. This indicates that low concentrations of TPC2-A1-N evoke a localised release of Ca^2+^ which is insufficient to evoke CICR unless contextually localised to IP_3_Rs. When cells were stimulated with histamine in the presence of TPC2-A1-N, the response to histamine was exaggerated. The sensitized histamine-evoked Ca^2+^ response prevented clustering of Rab46 at the MTOC and increased Ang2 secretion further implying TPC2 as a prominent player in this pathway. Importantly, the activity of TPC2 is specific to this pathway because whilst thrombin evokes secretion of Ang2 we know that thrombin does not activate Rab46-dependent trafficking of WPBs to the MTOC [8] nor does it evoke an NAADP-sensitive signal [31]. Similarly, global Ca^2+^ rises initiated from a plasma membrane cation channel do not evoke secretion of Ang2 nor trafficking of Rab46 and WPBs.

In total, this signalling pathway allows selective extracellular stimuli to be uniquely coupled to a downstream physiological response. Whilst the GTPase activity of Rab46 permits trafficking of a subpopulation of WPBs to a unique locale, this study suggests TPC1/2 channels regulate the critical Ca^2+^ signal needed for these WPBs to escape and to secrete their cargo. Rab46 acts as the downstream effector of TPC1/2 activity and regulates WPB exocytosis and Ang2 secretion, the molecular switches that allow this anterograde trafficking are unknown. Moreover, Rab46, may also be part of the molecular machinery necessary for formation of this unique nanodomain, since a localised Ca^2+^ signal imparted by TPC2-A1-N can be promoted to a global signal upon Rab46 activation, this will require further exploration using GCAMP-tagged channels, high resolution imaging of organelles and genetic manipulation of Rab46.

Gaining insight into this cellular pathway is important due to the proangiogenic function of Ang2 and may reveal how deviations in these mechanisms contribute to Ang2-dependent progression of vascular diseases, thereby aiding new therapeutic strategies. Evidence regarding the role of TPCs in the vasculature is increasing and several studies implicate TPCs in angiogenesis [50] [40]. Moreover, TPCs contribute to vascular-based diseases such as hypoxia-induced pulmonary hypertension [51] and macular degeneration [52]. In agreement Rab46 knockdown impacts the formation of blood vessels [11] and in human studies Rab46 SNPs have been associated with diseases associated with a dysfunctional endothelium such as COVID [53] and NAFLD [54, 55], in addition to immunodeficiencies [56] [57].

Limitations of this study: There could be off target effects of the pharmacological agents. To mitigate, we used chemically independent antagonists (Ned19 and tetrandrine) and tested an agonist (TPC2-A1-N) for opposing effects, which occurred.

## Materials and Methods

### Cell Culture

Pooled human umbilical vein endothelial cells (HUVECs: PromoCell) cultured in endothelial cell basal medium (EBM-2) supplemented with endothelial growth medium-2 supplement pack (EGM-2, PromoCell). Cells were incubated in a humidified incubator at 37°C and 5% CO_2_ and used between P1 - 5 for experimental purposes.

Primary human aortic endothelial cells (hAECs: PromoCell) were cultured in endothelial cell basal medium (EBM-2) supplemented with endothelial medium supplement pack MV2 (EMV-2, PromoCell). Cells were incubated in a humidified incubator at 37°C and 5% CO_2_ and used between P1 - 5 for experimental purposes.

### Intracellular Ca^2+^ Measurements – Flexstation

hAECs seeded on to a clear-bottom black Greiner Bio-One^®^ 96-well plate at a density of 200 µl max vol./15,000 cells per well and incubated at 37°C, 5% CO_2_ for 24 hrs to achieve a confluent monolayer. Recordings were undertaken using 1.5 mM Ca^2+^ SBS, or 0 mM Ca^2+^ SBS as stated. 24 hours after plating hAECs were loaded with 50 μl of Fura-2-AM solution ([1:500] Fura-2 (1 mM), 10% pluronic acid (PA-127), 1.5 mM Ca^2+^ SBS) for 1 hr at 37°C in the dark. Cells washed 2x with 100 µL SBS, the 2^nd^ wash was left on (specified inhibitors applied at this time point) and incubated for 30 mins at room temperature to allow cleavage of the AM ester. SBS was aspirated and 80-100 µL/ well recording buffer (SBS) added. The Flex station protocols were set up as described in the graphs and figure legends.

### Immunofluorescent staining

50,000 hAECs in 300 μL EMV-2 were incubated in an Ibidi^®^ µ-Slide 8-well^HIGH^ slide for 48 hrs (media change at 24 hrs) at 37°C, 5% CO_2_ in a humidified incubator. After 48 hrs cells were starved using 300 µL of M199 / well for 1 hr prior to stimulation. Inhibitors were added 30 mins into this starvation stage in M199 at the stated concentration. Cells were washed 1x with PBS and stimulated with histamine, Yoda1, TPC2-A1-N/P or vehicle control at the stated concentrations for 10 mins prior to fixation with 4% paraformaldehyde for 10 mins. Wells washed 3x with PBS and permeabilised with 0.1% Triton 10 mins. Cells were incubated with respective primary antibodies (1°) (see Table 2) at required dilution for 1 hr at room temperature. Unbound 1° antibody was removed with 3x PBS washes and cells incubated with fluorescently labelled Alexa Fluor secondary antibodies (2°) (see Table 2) at desired concentration for 30 mins in dark conditions. Cells washed 3x with PBS to remove unbound 2° antibody and the nucleus was stained using Hoechst 33342 at [1:1000 PBS] 7 mins. Wells were mounted with 4 drops of Invitrogen^®^ Prolong Gold mountant.

**Table 1:**
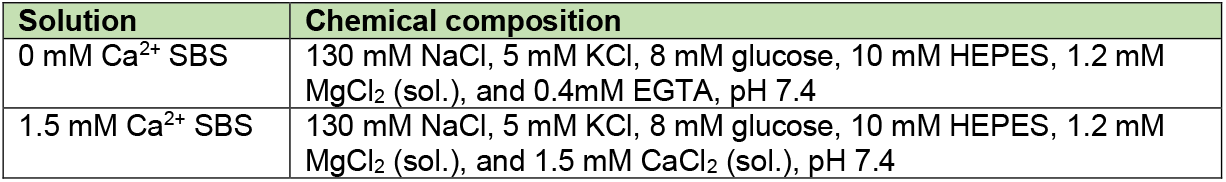
Buffers used for Ca^2+^ imaging.

**Table 2:** Antibodies use in immunofluorescent imaging.

| Primary antibody | Species | Dilution | Manufacturer | Lot. number |
| --- | --- | --- | --- | --- |
| Anti-vWF | Human | 1:200 | Abcam | AB778 |
| Anti-CRACR2A | Rabbit | 1:300 | Antibodies.com | A89955 |
| Anti-Pericentrin | Rabbit | 1:300 | Abcam | AB28144 |

| Secondary antibody | Species | Dilution | Manufacturer | Lot. number |
| --- | --- | --- | --- | --- |
| Alexa Fluor 488 Anti-Rabbit Fab Frag | Rabbit | 1:300 | Jackson ImmunoResearch Labs | 711-547-003 |
| Alexa Fluor 594 Anti-Mouse IgG | Mouse | 1:300 | Jackson ImmunoResearch Labs | 715-585-150 |

### Angiopoietin-2 ELISA

hAECs were seeded in Nunc® coated 96-well plate (Thermo Scientific) at a concentration of 15,000 cells/well, in total volume of 200 µL EMV2. After 24 hrs, the cells were incubated in basal M199 for 1 hr (+/- inhibitor) before stimulating with defined compounds for 10 mins in 1.5 mM Ca^2+^ SBS. The supernatant was collected and centrifuged at 500 x g for 5 mins and used immediately. The Human Angiopoietin-2 ELISA Kit (Antibodies.com, A77682) was used according to the manufacturer’s instruction except the use of non-diluted samples. Ang2 concentrations were calculated according to the generated standard curve. All raw data was exported to Microsoft Excel before final analysis was completed on GraphPad Prism 9 and expressed as % of control.

### Widefield deconvolution microscopy

The CellSens deconvolution system (Olympus) was used throughout using an Olympus IX-83 inverted microscope. For the purposes of Z-stack imaging, 10-focal planes where sectioned at 0.2 µm per Z-stack per image and were taken on a Photometrics-BSI camera device. Re-iterative deconvolution (5x) was performed using an advanced maximum likelihood algorithm to rebalance out of focus light (deconvolution) from one focal plane of the Z-stack to another. The filters used on the Olympus IX83 camera were: DAPI; GFP, and RFP. The objectives used were 60x/1.4 oil. All images were taken at ambient room temperature and image acquisition and furthering processing was conducted using the CellSens Olympus software. Images were acquired in a non-bias fashion using the computer module to move around the centre of the well, 5x images / condition. Exposure times for all images within each biological repeat were set equally for each filter according to the positive control (cells treated with 30 μM His).

### Image processing and analysis

All maximum projection (intensity) images were generated using the CellSens (Olympus) software before being transferred to ImageJ Fiji for analysis. ImageJ macros (see appendix 1) were used for the quantification of Rab46 distribution and WPB counts. **WPB counts:** Image channels were split into individual-coloured channels, and the background (3 px, sliding paraboloid) subtracted. A local threshold was set and subsequently followed by particle analysis (WPB count macro, see appendix 1) to determine the number of WPBs per image and per cell. Five regions of interest per condition were analysed and with three biological repeats. **Rab46 distribution:** The intracellular distribution of Rab46 and the particle intensity was determined as previous [8], using a customised macro (Rab46 distribution macro, see appendix 1) Briefly, an 8-bit image was loaded into ImageJ Fiji. Image channels were then split, and the background subtracted from the DAPI/GFP channels. A binary mask of the DAPI channel was created using a default threshold. Background noise was subtracted using the median filter and a distance map was generated using the StarDist 2D plugin (https://imagej.net/plugins/stardist) to estimate the cell edges. Each channel was segmented using the Max Entropy threshold and the distance and intensity of each particle from the nucleus was measured. The macro automatically exports the results tables with distance (Min) and intensity values per particle (IntDen) to an Excel file for each image, which are imported to GraphPad Prism 9 for analysis. Distance (Min) values ranged from 0 px (nucleus) to 255 px (periphery). This spectrum of values was then binned into 3 distinct regions: Perinuclear (x < 2 µm); Intermediate (2 µm < > 5 µm); and Periphery (x > 5 µm), where x is defined as the distance from the nucleus to the cells peripheral edge. The integrated intensity was then normalised to the total fluorescence of the image (TIF). The mean values were calculated for each area and averaged amongst all the images per condition. The mean values amongst all biological repeats were presented in GraphPad Prism 9 as bar-plots depicting mean ± SEM, whereby the Y-axis denotes the ‘normalised integrated intensity (as ratio of TIF) and the X-axis denoting the distance from nucleus displayed as binned signal into the three subcellular regions described above.

### Data Analysis

All averaged data is presented as the mean SEM. Any outliers were detected and then subsequently removed using a Grubb’s test and equal variance established. To determine if the data was extracted from a normally distributed population, a Shapiro-Wilk normality test was performed. When comparing data from two groups (normally distributed) significance was assessed using a two-sample t-test or a one-way or two-way ANOVA coupled with the Tukey *post-hoc* test when considering three or more groups of data. The test chosen was dependent on the number of groups being considered. The presence of statistical significance was determined to exist at probability *p* < 0.05 (**** < 0.0001, *** < 0.001, ** < 0.01, * < 0.05). ‘ns’ signifies no significant differences between groups were observed. GraphPad PRSIM9 and OriginPro22b were used for computing data analysis as well as presentation of analysed data. *n* = the number of independent biological repeats, N = total number of technical repeats per condition. For immunofluorescence microscopy, each technical repeat of each condition included 5-10 individual images. Fitted Ca^2+^ concentration response curves were plotted using a Hill Equation indicating the 50 % maximum effect.

## Acknowledgements

This research was supported by a BHF PhD studentship (FS/PhD/21/29176) awarded to Ryan Murray and a BHF PhD studentship as part of a 4yr BHF programme awarded to Melissa Rose (FS/4yrPhD/F/24/34209). We’d like to thank the University of Leeds, Faculty of Biological Sciences Bioimaging facility for their advice (https://biologicalsciences.leeds.ac.uk/facilities/doc/bio-imaging-flow-cytometry).

## Author Contributions

RDM, MR and KTM performed the experiments. RDM, DJB and LM analysed the data and created the tables. LM conceived the study and co-wrote the article with input from all authors. LM generated research funds and supervised the PhD studentships.

