## Supplementary data for "Two-pore channel protein 2-mediated calcium release promotes angiopoietin 2 secretion by regulating Rab46-dependent Weibel-Palade body trafficking"

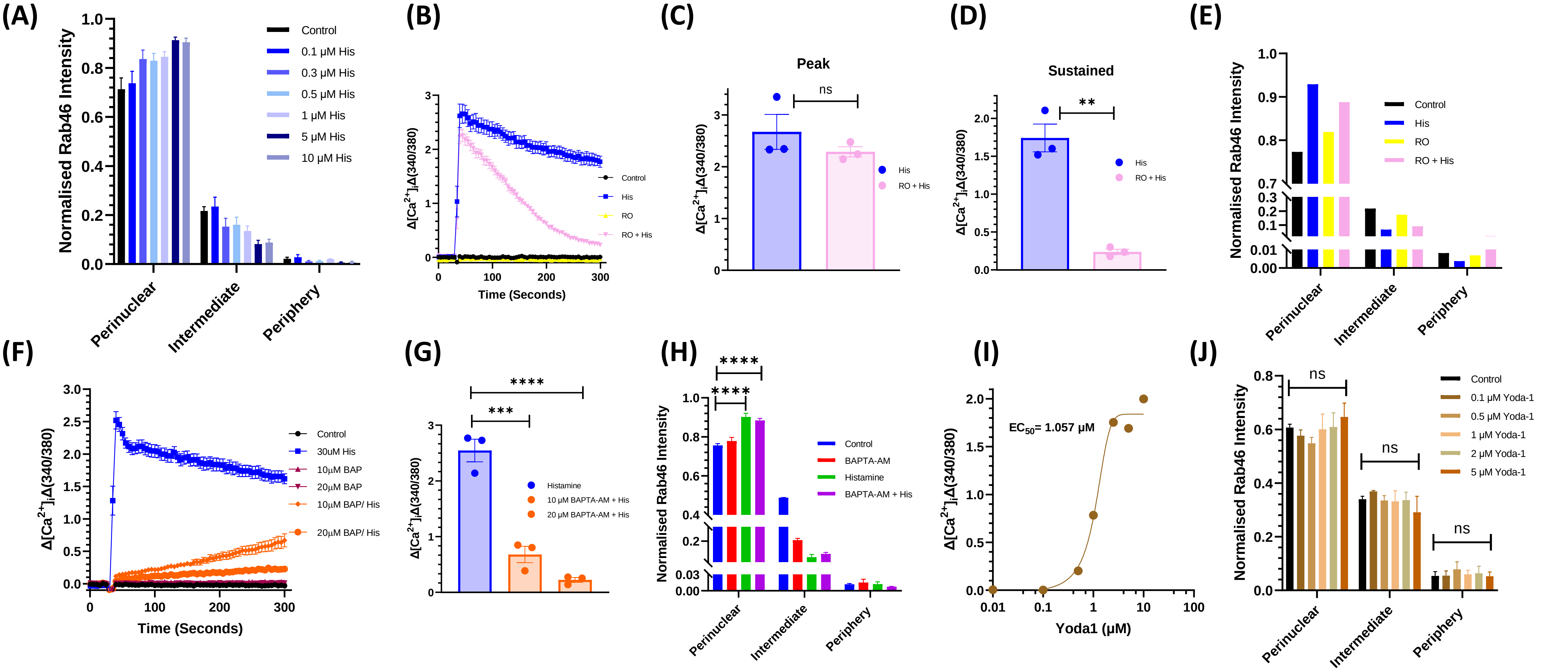


**Supplementary Fig. 1**. (A) Quantitative analysis of Rab46 cellular distribution in hAECs upon increasing [histamine]. Results were grouped into three areas as per methods. The plot shows Rab46 signal intensity of each particle in the respective area where the mean (± SEM) was noted as percentage of the total signal intensity. (B – D). Representative traces (B) and mean peak (C) / sustained (D) responses measuring changes (Δ) in intracellular $Ca^{2+}$ evoked by 30 μM histamine in the presence or absence of 10 μM of the Orai1 channel inhibitor RO5929. (E) Quantitative analysis of Rab46 cellular distribution in hAECs pre-treated with RO5929 before stimulation with histamine or control as depicted in images (Fig. 1C). Results were grouped into three areas and analysed as described above. (F - H) Representative traces (F) and mean peak (G) responses measuring changes (Δ) in intracellular $Ca^{2+}$ evoked by histamine in the presence or absence of 10 μM or 20 μM of the fast $Ca^{2+}$ chelator BAPTA-AM. (H) Quantitative analysis of Rab46 cellular distribution in hAECs depicted in images (Fig. 1E). Results were grouped into three areas and quantified as described above. (I) Mean data of hAECs treated with the Yoda1, (0.1 – 10 μM: an agonist of the plasma membrane $Ca^{2+}$ channel Piezo1) displayed as a concentration-response curve.  n/N = 3/9, error bars represent SEM. (J) Quantitative analysis of Rab46 cellular distribution in hAECs upon increasing Yoda1 stimulation. For each cellular distribution analysis n/N=3/15. *p-value < 0.05 from one-way ANOVA with Tukey post-hoc and unpaired t-tests.

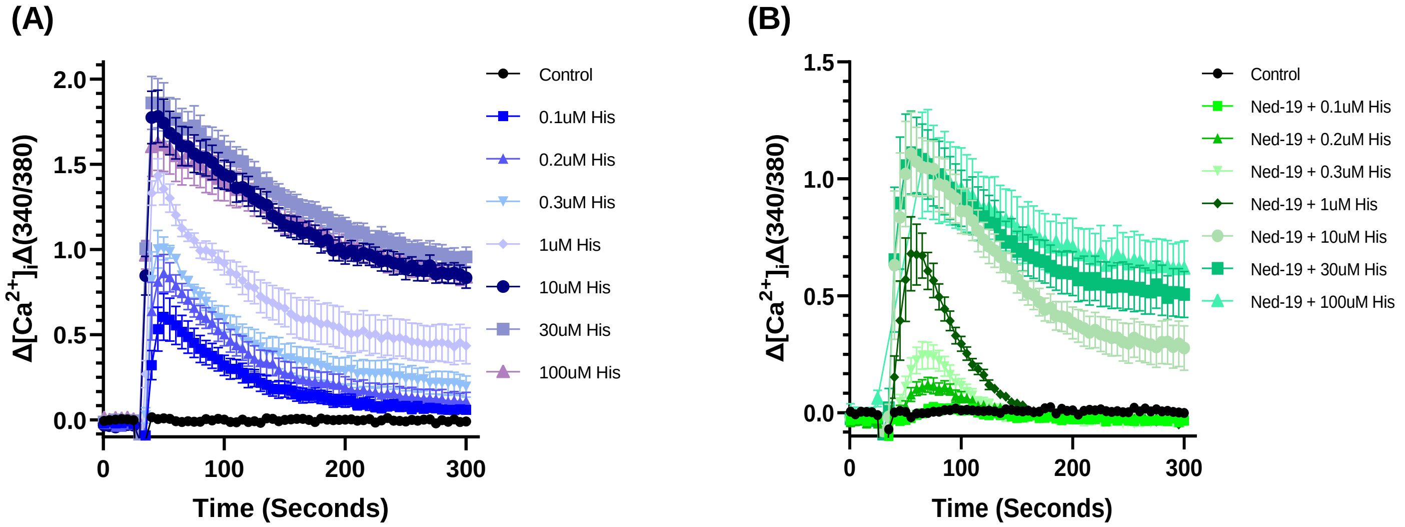


**Supplementary Fig. 2**. Representative traces measuring changes (Δ) in intracellular $Ca^{2+}$ evoked by increasing concentrations of histamine preincubated with vehicle control (A) or 100 μM Ned19 (B).

**(A)**

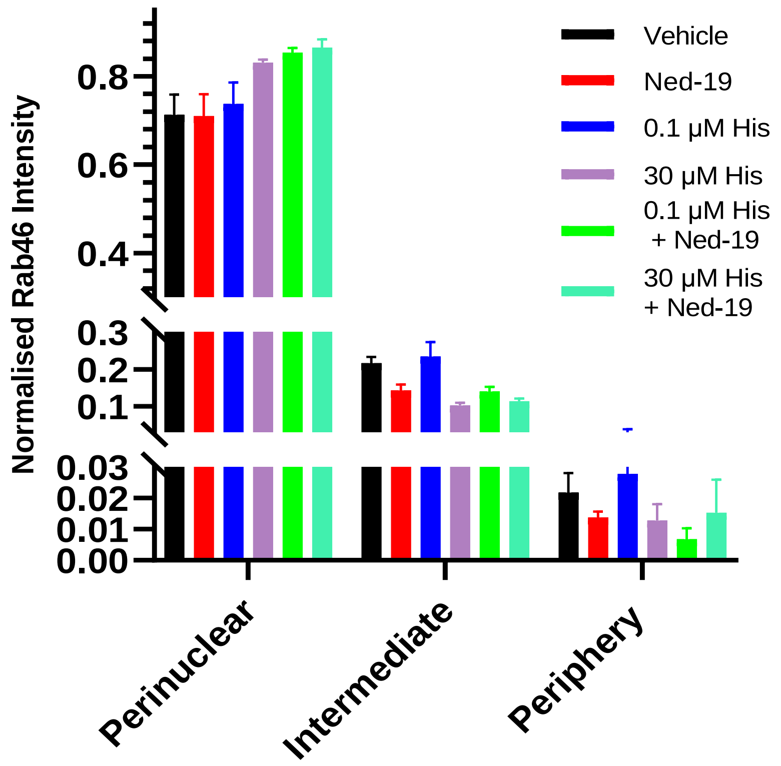


**Supplementary Fig. 3.** (A) Quantitative analysis of Rab46 cellular distribution in hAECs stimulated with increasing concentrations of histamine in cells preincubated with vehicle control or 100 µM Ned19. n/N=3/15.

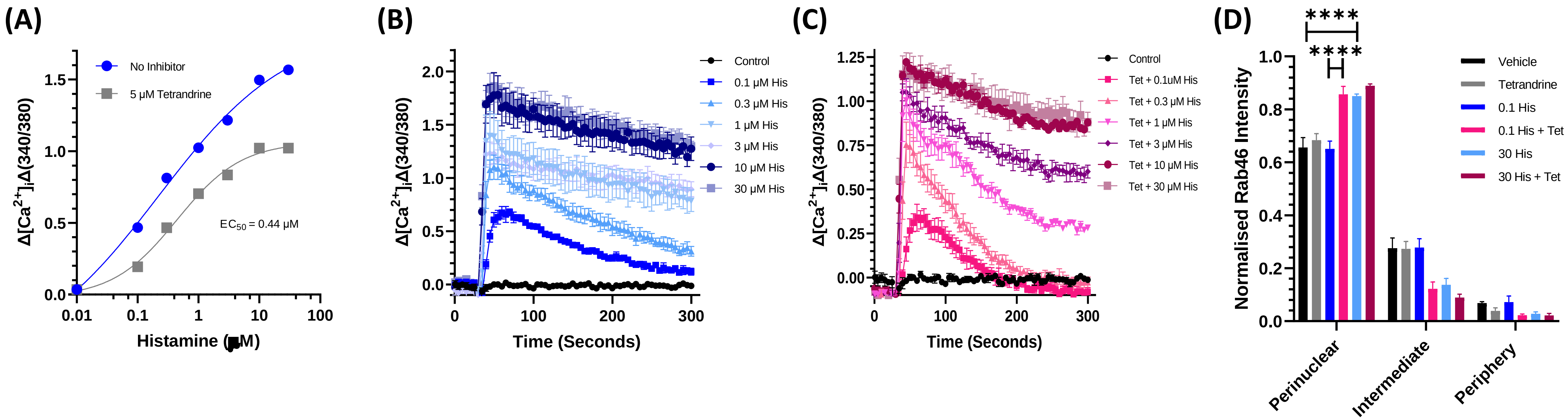


**Supplementary Fig. 4.** (A) Mean data of hAECs treated with tetrandrine displayed as a concentration-response curve. n/N = 3/9. Representative traces measuring changes (Δ) in intracellular $Ca^{2+}$ evoked by increasing concentrations of histamine preincubated with vehicle control (B) or 100 μM tetrandrine (C). (D) Quantitative analysis of Rab46 cellular distribution in hAECs stimulated with histamine in cells preincubated with vehicle control or 5 μM tetrandrine. Results were grouped into three areas: perinuclear, intermediate and periphery as described in methods. The plot shows Rab46 signal intensity of each particle in the respective area where the mean (± SEM) was noted as percentage of the total signal intensity. n/N=3/15. *p-value < 0.05 from 2-way ANOVA with Tukey post hoc analysis.

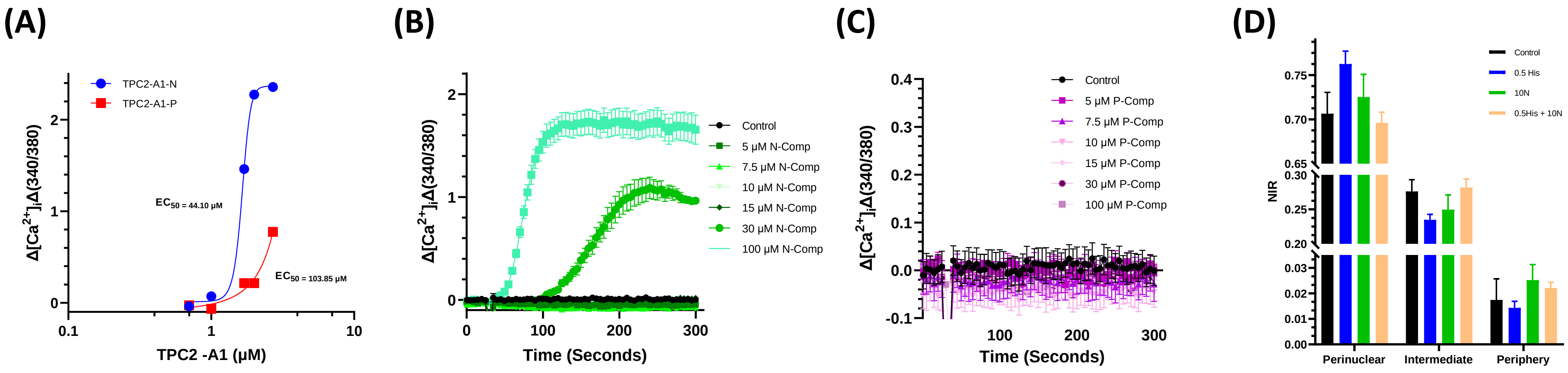


**Supplementary Fig. 5. (A)** Mean data of HUVECs treated with (0.1-100 μM) TPC2-A1-N or TPC2-A1-P displayed as concentration-response curves. n/N = 3/9. (B) Representative traces measuring changes (Δ) in intracellular $Ca^{2+}$ evoked by increasing concentrations of TPC2-A1-N in hAECs. (C) Representative traces measuring changes (Δ) in intracellular $Ca^{2+}$ evoked by increasing concentrations of TPC2-A1-P in hAECs. (D) Mean cellular distribution analysis of Rab46 in hAECs treated with 0.5 μM histamine alone or simultaneous addition of 0.5 μM histamine and 10 μM TPC2-A1-N. Results were grouped into three areas: perinuclear, intermediate and periphery as described in methods. The plot shows Rab46 signal intensity of each particle in the respective area where the mean (± SEM) was noted as percentage of the total signal intensity. n/N=3/15. *p-value < 0.05 from 2-way ANOVA with Tukey post hoc analysis.

**Supplementary Appendix 1**

**Weibel-Palade Body distribution macro for ImageJ**

```
imageTitle=getTitle();//returns a string with the image title

run("Split Channels");

selectWindow("C1-"+imageTitle);

close();

selectWindow("C2-"+imageTitle);

run("Subtract Background...", "rolling=3 sliding");

run("Duplicate...", " ");

rename("DUPLICATE");

selectWindow("C3-"+imageTitle);

rename("nuclei");

run("Command From Macro", "command=[de.csbdresden.stardist.StarDist2D],
args=['input':'nuclei', 'modelChoice':'Versatile (fluorescent nuclei)', 'normalizeInput':'true',
'percentileBottom':'1.0', 'percentileTop':'99.8', 'probThresh':'0.5', 'nmsThresh':'0.4',
'outputType':'Both', 'nTiles':'1', 'excludeBoundary':'2', 'roiPosition':'Automatic', 'verbose':'false',
'showCsbdeepProgress':'false', 'showProbAndDist':'false'], process=[false]");

selectWindow("Label Image");

setOption("ScaleConversions", true);

run("8-bit");

run("Auto Local Threshold", "method=Contrast radius=15 parameter_1=0 parameter_2=0
white");

setOption("BlackBackground", true);

run("Convert to Mask");

run("Invert LUTs");

run("Distance Map");

run("Invert LUTs");

selectWindow("C2-" +imageTitle);

//run("Brightness/Contrast...");

setMinAndMax(60, 255);

run("Apply LUT");

selectWindow("C2-" +imageTitle);

run("Auto Threshold", "method=MaxEntropy white");
```

```
run("Set Measurements...", "area mean min integrated redirect=[Label Image] decimal=3");
selectWindow("C2-" +imageTitle);
run("Analyze Particles...", "size=0.01-Infinity display");
saveAs("Results", "\\C:\\Users\\INSERT FILE PATH \\" + imageTitle + ".txt"); MAP
close("Results");
selectWindow("C2-"+imageTitle);
run("Set Measurements...", "area mean min integrated redirect=DUPLICATE decimal=3");
selectWindow("C2-"+imageTitle);
run("Analyze Particles...", "size=0.01-Infinity display");
saveAs("Results", "\\C:\\Users\\INSERT FILE PATH \\" + imageTitle + ".txt"); C2
close();
close("Results");
close("ROI Manager");
close("DUPLICATE");
close();
close();
```
